# Mapping chromatin state and transcriptional response in CIC-DUX4 undifferentiated round cell sarcoma

**DOI:** 10.1101/2023.10.11.561932

**Authors:** Nicholas J. Thomas, Cuyler Luck, Nicole Shlimon, Rovingaile Kriska Ponce, Zeinab Kosibaty, Ross A. Okimoto

**Author notes:** Correspondence to: Ross A. Okimoto, University of California, San Francisco 513 Parnassus Avenue, HSW1201P San Francisco, CA 94143. Equal contribution (Nicholas J. Thomas, Cuyler Luck). Conflict of interest: The authors declare no potential conflicts of interest.

## Abstract

CIC-DUX4 is a rare and understudied transcription factor fusion oncoprotein. CIC-DUX4 co-opts native gene targets to drive a lethal form of human sarcoma. The molecular underpinnings that lead to oncogenic reprograming and CIC-DUX4 sarcomagenesis remain largely undefined. Through an integrative ChIP and RNA-Seq analysis using patient-derived CIC-DUX4 cells, we define CIC-DUX4 mediated chromatin states and function. We show that CIC-DUX4 primarily localizes to proximal and distal cis-regulatory elements where it associates with active histone marks. Our findings nominate key signaling pathways and molecular targets that enable CIC-DUX4 to mediate tumor cell survival. Collectively, our data demonstrate how the CIC-DUX4 fusion oncoprotein impacts chromatin state and transcriptional responses to drive an oncogenic program in undifferentiated sarcoma.

**Significance:** CIC-DUX4 sarcoma is a rare and lethal sarcoma that affects children, adolescent young adults, and adults. CIC-DUX4 sarcoma is associated with rapid metastatic dissemination and relative insensitivity to chemotherapy. There are no current standard-of-care therapies for CIC-DUX4 sarcoma leading to universally poor outcomes for patients. Through a deep mechanistic understanding of how the CIC-DUX4 fusion oncoprotein reprograms chromatin state and function, we aim to improve outcomes for CIC-DUX4 patients.

## Introduction

Transcription factor (TF) fusion oncoproteins represent a prototypical example of how a key genetic alteration can drive cellular transformation and oncogenesis (1,2). Unlike the majority of human cancers that are associated with multiple genetic and non-genetic changes, TF fusion oncoproteins are often observed in relatively “silent” cancer genomes (3–5). These findings indicate that TF fusions may constitute a disease-defining genetic event that underlies cancer pathogenesis.

One key example is the CIC-DUX4 TF fusion that results from a chromosomal translocation involving either t(4;19) or t(10;19). CIC-DUX4 sarcomas are the most common genetic alteration detected in *EWSR1*-negative small round cell sarcomas (6,7). CIC-DUX4 rearrangements are associated with highly aggressive and lethal clinical phenotypes, including a high metastatic rate, chemo-insensitivity, and poor overall survival (6). Furthermore, there is no consensus standard-of-care treatment option for CIC-DUX4 patients, who often receive ineffective salvage chemotherapy.

Consistent with other TF fusion oncoproteins (3,4), CIC-DUX4 rearrangements are associated with silent cancer genomes yet it constitutes a disease-defining molecular driver of sarcomagenesis (5). These observations provide a clean genetic background to better understand the regulatory mechanisms by which fusions, including CIC-DUX4 lead to target gene activation and/or repression. Thus, while we and others have previously reported CIC-DUX4 binding and CIC-DUX4 mediated gene expression patterns (8–10), the key chromatin remodeling events associated with gene activation/repression remain relatively unknown. To meet this need, we integrated measurement of CIC-DUX4 binding and key histone marks via ChIP-Seq with RNA-seq gene expression changes in patient-derived cells to systematically analyze the chromatin state and function of endogenous CIC-DUX4.

## Results

### CIC-DUX4 binds to poised and active chromatin states through cis-regulatory elements

The human cancer genome is typically divided into active (open chromatin) euchromatin and repressed (closed chromatin) heterochromatin. Active euchromatic regions are marked by histone modifications that enable chromatin accessibility and TF binding, leading to gene transcription (11). In contrast, repressed heterochromatin is associated with repressive histone marks that decrease TF accessibility (11). Using well-defined histone marks (11,12) that correlate with active (H3K27ac – active enhancer, H3K4me1 – primed enhancer, H3K4me3 – primed promoter) or repressed (H3K27me3) chromatin states (13–15) we mapped the chromatin state of CIC-DUX4 sarcomas.

We first performed ChIP-Seq using a validated CIC-directed antibody (9) to identify endogenous CIC-DUX4 binding sites in a patient-derived CIC-DUX4 sarcoma cell line (NCC-CDS1-X1-C1) (Figure 1A). Through Model-based Analysis of ChIP-Seq (MACS) and focusing on peaks located within 1kb of an activating histone mark (H3K27ac, H3K4me3), we identified 392 high-confidence (FDR q<0.001) CIC-DUX4 peaks. Consistent with prior work, the majority of CIC-DUX4 binding occurred in intergenic (44%) and intronic regions (50%), with a small fraction (3%) mapping to promoters in NCC-CDS1-X1-C1 cells (Figure 1B, Supplemental Figure 1A-B). In order to associate these CIC-DUX4 binding sites with regulatory elements, we evaluated signals of the aforementioned key histone marks (H3K27ac, H3K4me1, H3K4me3, and H3K27me3) (16) at these high-confidence CIC-DUX4 peaks (Figure 1A). We observed that the majority of CIC-DUX4 peaks correlated with H3K4me1 sites, a marker associated with poised cis-regulatory elements (13). To correlate CIC-DUX4 peaks with active sites of transcription we mapped H3K27ac (enhancers) and H3K4me3 (promoters) marks and compared them to CIC-DUX4 binding. We noted that H3K27ac and H3K4me3 marks were associated with a fraction of CIC-DUX4 binding sites, suggesting that CIC-DUX4 can indeed activate transcription through cis-regulatory elements (Figure 1C-D).

**Figure 1:**
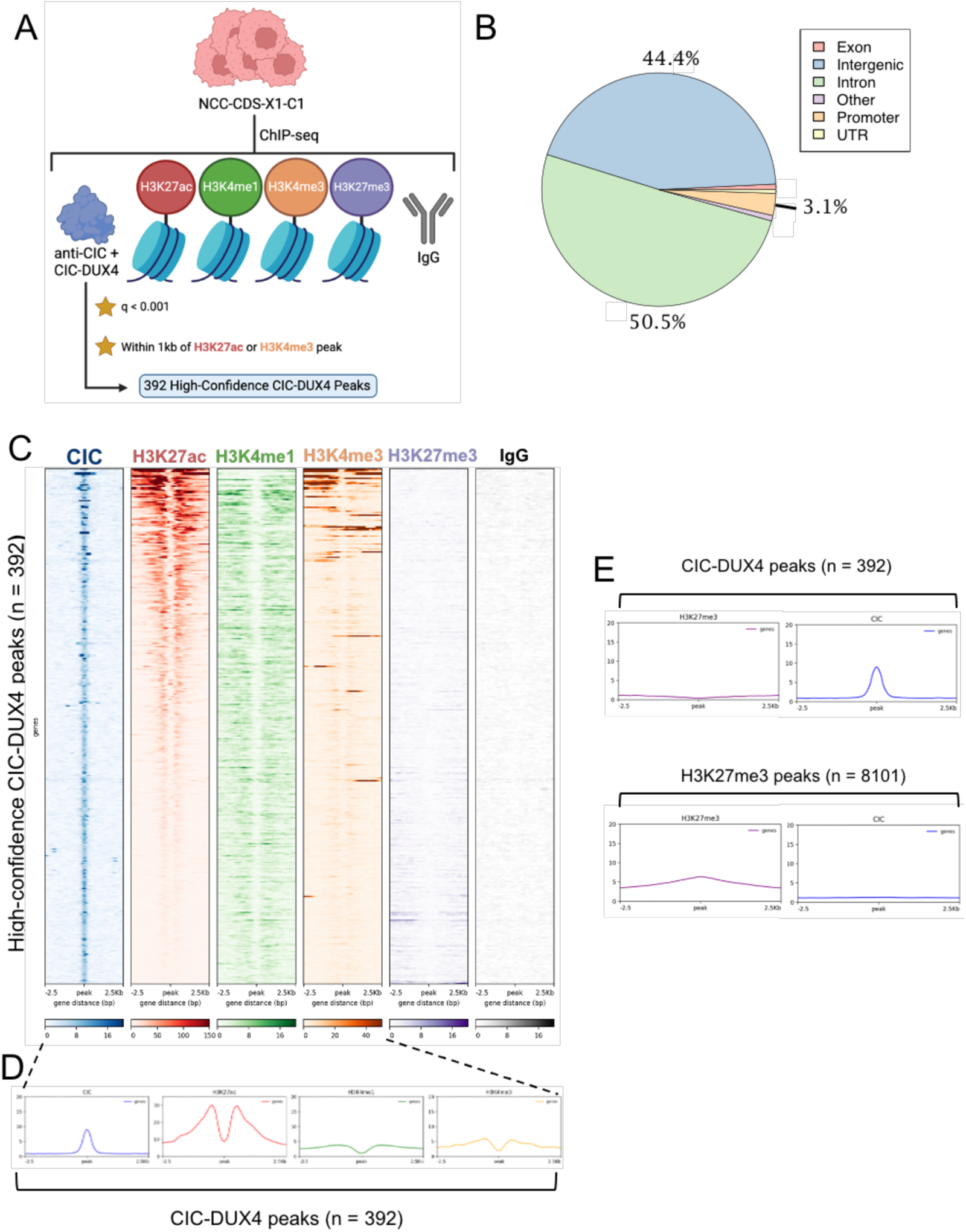
High-confidence CIC-DUX4 ChIP-seq peaks associate with activating histone marks. A) Schematic algorithm of ChIP-seq and identification of the 392 high-confidence CIC-DUX4 peaks in NCC-CDS-X1-C1 cells. Image generated with BioRender (XJ25WRKTK3). B) Distribution of genomic elements annotated for high-confidence CIC-DUX4 peaks. C) deeptools heatmap of ChIP signal at all 392 high-confidence CIC-DUX4 peaks (rows) for all six ChIP targets, ordered by decreasing mean H3K27ac signal. D) deeptools profile plots from data analyzed in C. E) deeptools profile plots of H3K27me3 and CIC signal at 392 high-confidence CIC-DUX4 peaks (data from C) or at 8101 H3K27me3 peaks (q < 0.001, peak in non-blacklisted region).

Since other TF fusions have been shown to regulate gene expression in both a positive and negative fashion, we wanted to determine if CIC-DUX4 was also associated with target gene silencing. To explore this, we performed H3K27me3 (modification by the polycomb repressive complex) ChIP-Seq to identify repressive sites in NCC-CDS-X1-C1 cells. We identified 8,101 H3K27me3 peaks but did not observe a significant overlap with CIC-DUX4 peaks (n = 392) (Figure 1E). These findings suggest that the CIC-DUX4 fusion oncoprotein largely interacts with chromatin at poised and active cis-regulatory elements.

### CIC-DUX4 mediated gene expression

In order to understand how CIC-DUX4 mediated chromatin remodeling impacts gene expression, we knocked down (KD) CIC-DUX4 expression using siRNA in two patient-derived CIC-DUX4 sarcoma cell lines (NCC-CDS-X1-C1 and NCC-CDS2-C1) (17,18) and made comparisons between siControl and siCIC (Figure 2A and Supplemental figure 2A-B). We confirmed the on-target specificity of our CIC-DUX4 siRNA through quantification of a 20bp trans-breakpoint sequence (breakpoints span across CIC exon 20 and DUX4 exon 1, with slight differences for each cell line) within CIC-DUX4 (Figure 2B, bottom panel). This reduction in trans-breakpoint reads could not be explained by differences in total read counts, implying successful gene silencing (Figure 2B, top panel). Genetic CIC-DUX4 suppression resulted in 1080 and 179 downregulated and 2718 and 207 upregulated genes in NCC-CDS-X1-C1 and NCC-CDS2-C1, respectively (Figure 2C, q-value cutoff 0.0001, log2FC cutoff +/-0.58). Our CIC-DUX4 mediated gene expression data is consistent with other prior work that identified high-confidence CIC-DUX4 target genes in independent CIC-DUX4 model systems (e.g. IB120) (19). Specifically, we observed an overlap between several of these 25 high-confidence CIC-DUX4 target genes (yellow data points) and the significantly downregulated genes in NCC_CDS1_X1_C1 expressing CIC siRNA. These findings suggest that a highly conserved CIC-DUX4 mediated transcriptional program may drive sarcomagenesis. To further understand the key pathways that CIC-DUX4 regulates to promote tumor progression, we performed gene ontology (GO) using the DAVID (20) and PANTHER (21) tools. We found that CIC-DUX4 mediates multiple pathways associated with cell cycle and DNA replication and repair, which is consistent with prior studies (22). We next performed an integrative RNA-Seq and ChIP-Seq based analysis to identify both known and novel CIC-DUX4 target genes. Using our 1,125 high confidence RNA-seq and the 335 ChIP-seq derived gene annotations we identified 37 common genes including known (ie. *VGF, CCND2, DUSP4*) and as of yet uncharacterized molecular targets (ie. *SPRED2*).

**Figure 2:**
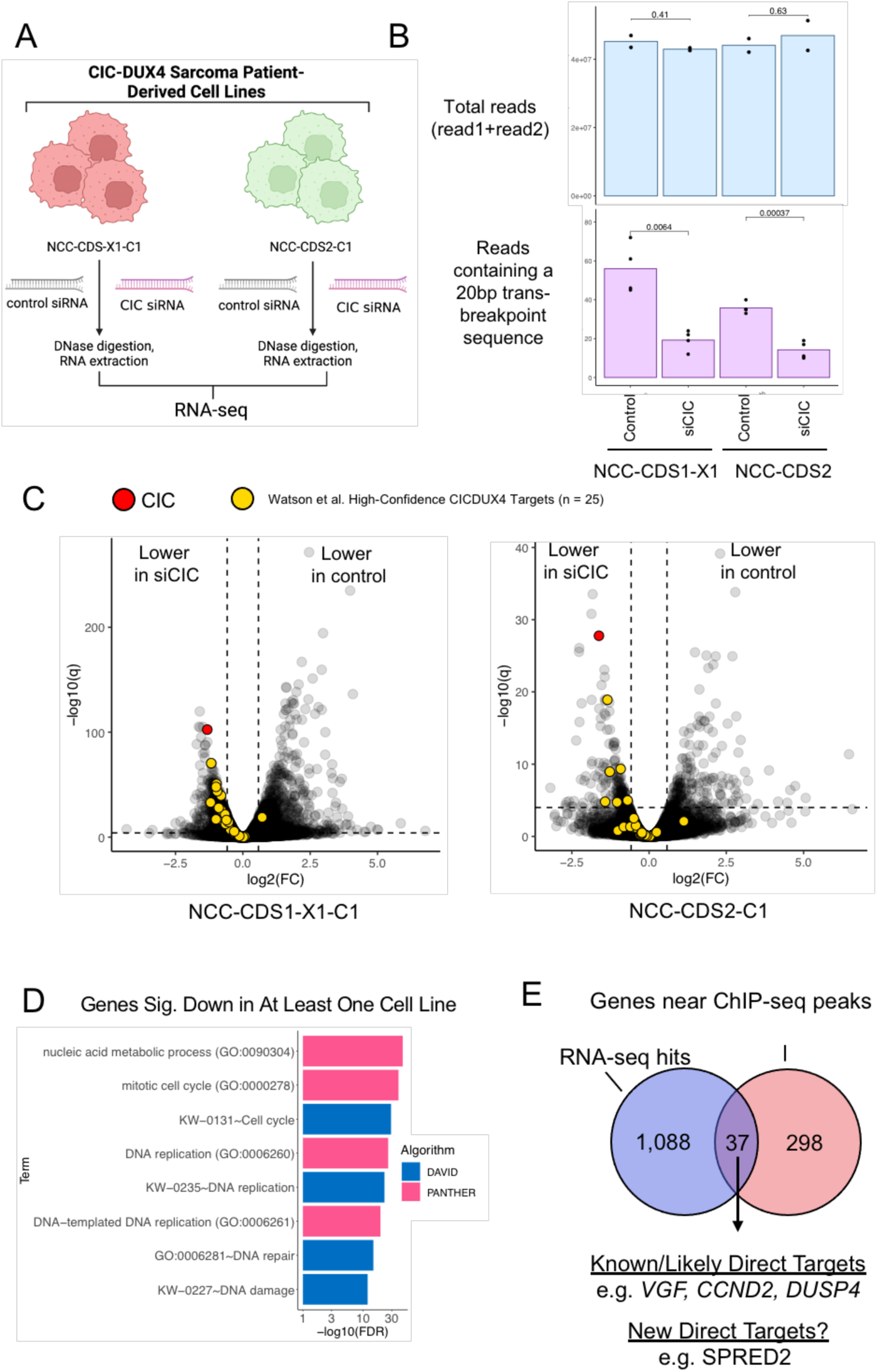
Identification of CIC-DUX4-mediated transcriptional targets in patient-derived cell lines. A) Schematic workflow to identify CIC-DUX4 regulated transcriptional targets in NCC-CDS-X1-C1 and NCC-CDS2-C1 cells. Image generated with BioRender (AO25WRL54B). B) Total reads (read1+read2) returned by Novogene for each sample (top) or reads from each read file filtered by using the “grep” command for a 20 base trans-breakpoint sequence (bottom). P-values shown were calculated by Student’s t-test. C) Volcano plots of genes processed with edgeR for NCC-CDS-X1-C1 and NCC-CDS2-C1 cells. Genes colored yellow correspond to the 25 high-confidence CIC-DUX4 target genes identified in (https://doi.org/10.1101/517722). Significance cutoffs displayed correspond to -log10(q) = 4, and |log2(FC)| > 0.58. D) Selected gene ontology terms from DAVID and PANTHER analyses, chosen for their significance and biological meaning. E) Venn diagram of genes significantly downregulated (q < 0.0001, log2FC < -0.58) in the RNAseq dataset for at least one cell line (left, n = 1125), and/or were annotated by HOMER as being the closest gene to a high-confidence ChIP-seq CIC-DUX4 peak (right, unique genes not repeatedly counted, n = 335).

### CIC-DUX4 binding at non-consensus TGNNTGNN motifs

The consensus CIC (Capicua) DNA-binding motif is T(G/C)AATG(G/A)A, which has been well defined in multiple drosophila models (23). While the wild-type CIC DNA-binding octamer is conserved in human species, it remains unclear if this is the preferred binding motif in the context of CIC-DUX4. To explore CIC-DUX4 binding more broadly, we performed motif-based analysis of our ChIP-Seq dataset using MEME-ChIP (24–26). Through this analysis we confirmed that the highly conserved wild-type CIC motif TGAATGAA was highly significant (Figure 3A). We further noted that positions 3&4 and 7&8 within the octameric sequence were variable (A or G) relative to the almost invariant TG sequence in positions 1&2 and 5&6 (i.e., TGNNTGNN). Thus, we hypothesized that CIC-DUX4 may potentially bind TGNNTGNN type motifs in NCC-CDS1-X1-C1 cells. To explore this further, we extracted all overlapping 8-mers from the sequences contained in the 374 unique genomic regions of the high-confidence CIC-DUX4 peaks (see Methods) and counted the number of times TGNNTGNN motifs appeared in each peak (Figure 3B). When we searched for TGNNTGNN motifs with the highest mean proportional frequency among all peak sequences, 11 TGNNTGNN variants were more common than the rest, including the consensus octamer TGAATGAA (Figure 3C). These variants largely used A/G nucleotides at positions 3/4 and 7/8, in line with our MEME-ChIP analysis. However, a given motif could occur at a high mean frequency either because it occurs in many sequences or it occurs many times in a low number of sequences. Therefore, we visualized how many times each of the 11 variants occurred in all 374 high-confidence peak regions (Figure 3D). Through this analysis we observed that nearly half of the sequences associated with CIC-DUX4 peaks contained none of the 11 most common TGNNTGNN variants. Of the remaining sequences, some contained few instances of single TGNNTGNN variants while several contained high numbers of many different TGNNTGNN variants. This led us to hypothesize that repetitive or redundant octameric variants may provide CIC-DUX4 binding sites beyond the canonical single- or double-TGAATGAA motifs.

**Figure 3:**
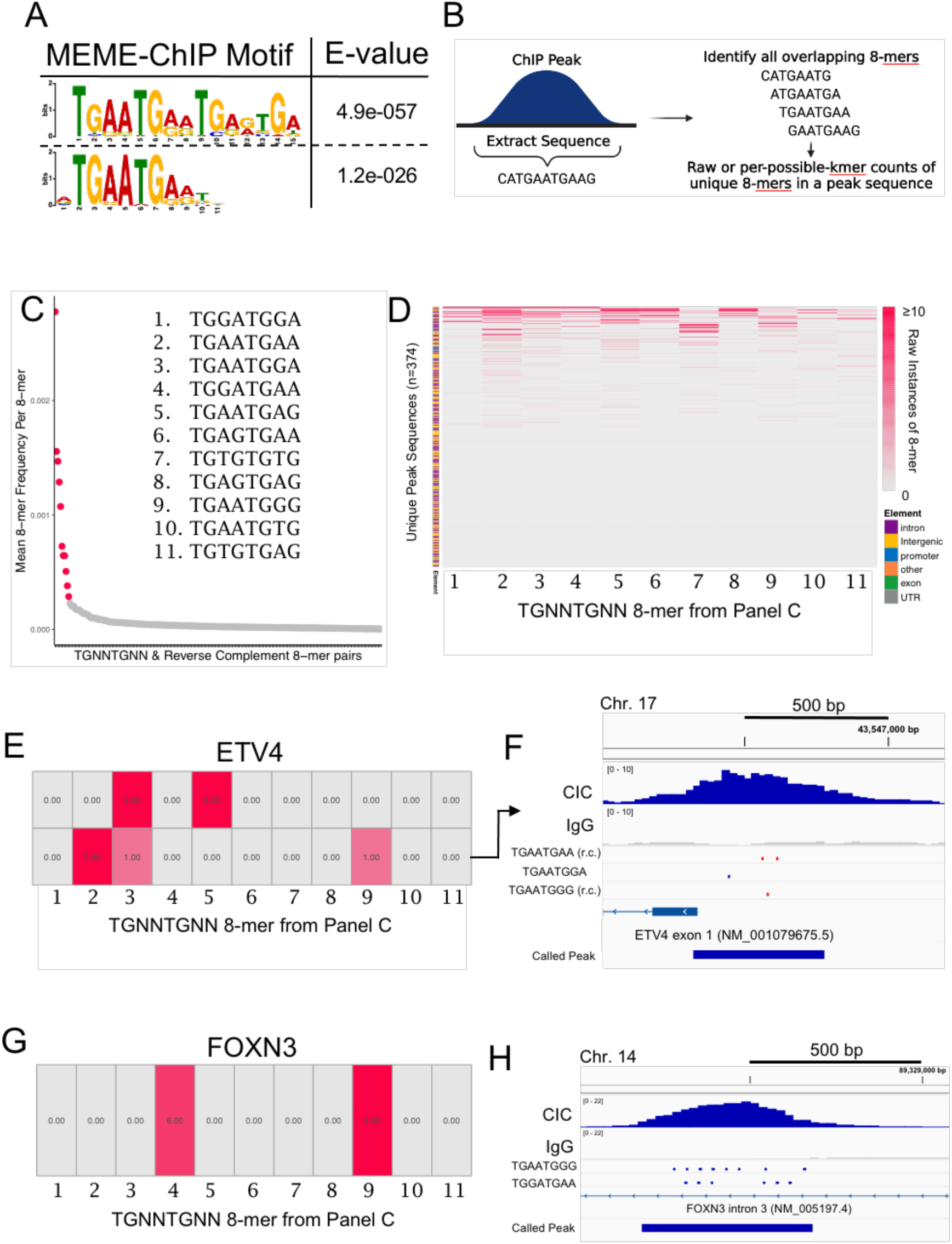
Repetitive non-canonical CIC binding motifs associate with CIC-DUX4 binding sites. A) MEME-ChIP analysis of the high-confidence CIC-DUX4 peaks confirmed enrichment of the known CIC octamer (TGAATGAA). B) Graphical representation of the analysis used to determine the frequency of 8-mers in the sequences underlying high-confidence CIC-DUX4 peaks (see Methods for details, Image generated in BioRender, VT25WRL6T0). C) Frequency of a given 8-mer divided by the number of total 8-mers in the sequence, averaged across all high-confidence CIC-DUX4 peaks. Inset: the sequences for the top eleven 8-mers ordered by descending mean normalized frequency. D) Heatmap of the raw occurrences of the eleven 8-mers from C (rows) in all unique CIC-DUX4 peak sequences (columns, n = 374). Color legend indicates the genomic element the peak was located in, as annotated with HOMER. E) Sub-section of D for the two CIC-DUX4 peaks annotated for *ETV4*. F) IGV visualization of the peak in the *ETV4* promoter, including the location of TGNNTGNN-variant sequences (r.c. = reverse complement). G) Sub-section of D for the CIC-DUX4 peak annotated for *FOXN3*. H) IGV visualization of the peak in the *FOXN3* intron, including the location of TGNNTGNN-variant sequences.

To further explore these octameric variants, we used *ETV4*, a known CIC-DUX4 target gene (27), and mapped a broad CIC-DUX4 peak on the *ETV4* promoter. While we noted tandem TGAATGAA motifs within this peak, we additionally identified two variant TGNNTGNN type repeats within the proximal regulatory element (TGAATGGA, TGAATGGG) (Figure 3E-F). Since CIC-DUX4 peaks primarily occur in intronic regions (50%), we next asked if CIC-DUX4 localized to TGNNTGNN sites within the introns of putative target genes. *FOXN3,* a TF that belongs to the forkhead family, harbors multiple repetitive TGAATGGG and TGGATGAA sites that correspond to a broad CIC-DUX4 peak on Chromosome 14 within intron 3 of the *FOXN3* gene (Figure 3G-H, Supplemental Figure 3A). POLE on chromosome 12 is another representative example of CIC-DUX4 binding at TGAATGAG repetitive octamer variant sites within an intron (Supplemental Figure 3B-C). Through the analysis of intergenic regions (44% of CIC-DUX4 peaks), we also identified repetitive TGNNTGNN motifs that are associated with CIC-DUX4 binding.

A representative example includes the CIC-DUX4 peak mapped to *CDH4*, which was the highest scoring peak in our dataset. This peak was associated with TGAATGGG repeats ∼9kb upstream of the transcriptional start site (Supplemental Figure 3D-E). These findings suggest that CIC-DUX4 are associated with and may potentially bind to TGNNTGNN variants, particularly repetitive elements. These findings help to explain how the majority of CIC-DUX4 peaks occur outside of promoter regions (3%) and/or at sites not associated with the consensus wild-type CIC octamer (T(G/A)AATG(G/A)A).

### Multi-level negative regulation of the MAPK signaling cascade by CIC-DUX4

We have previously identified that CIC-DUX4 directly binds and transcriptionally controls *DUSP6*, a negative regulator of the terminal MAPK substrate, ERK. In particular, CIC-DUX4 transcriptionally upregulates the ERK specific phosphatase, *DUSP6*, which inactivates ERK and augments CIC-DUX4 expression (active ERK leads to CIC-DUX4 degradation) (9). Thus, one interesting finding through this integrative analysis was the identification of other negative MAPK regulators, including multiple SPRED and SPROUTY family members that are known to silence more proximal MAPK substrates including Ras (28,29). These findings led us to hypothesize that CIC-DUX4 potentially regulates its own expression through multi-level feedback on the MAPK signaling cascade. Specifically, through direct regulation of *DUSP6* and potentially *SPRED/SPROUTY* family members, CIC-DUX4 auto-regulates its own expression. To test this, we first performed a Ras-GTP pulldown assay to assess Ras activity upon CIC-DUX4 expression. Active Ras (GTP-Ras) decreased upon exogenous CIC-DUX4 expression in 293T cells, suggesting that there is indeed a more proximal level of MAPK control by CIC-DUX4 (Figure 4A). To confirm CIC-DUX4 target gene regulation, we performed qRT-PCR on putative target genes including multiple SPRED and Sprouty family members that associated with CIC-DUX4 binding in our ChIP-Seq dataset (Figure 4B and Supplementary Figure 4A-D). SPRED1, which harbored a CIC-DUX4 ChIP-Seq peak within the proximal regulatory element (Figure 4B), increased upon exogenous CIC-DUX4 expression in 293T cells (Figure 4C-D) and decreased upon CIC-DUX4 KD in CIC-DUX4 patient derived cells (NCC-CDS-X1-C1) (Figure 4E). These findings suggest that CIC-DUX4 attenuates MAPK flux through direct transcriptional upregulation of negative MAPK-Ras-ERK regulators.

**Figure 4.**
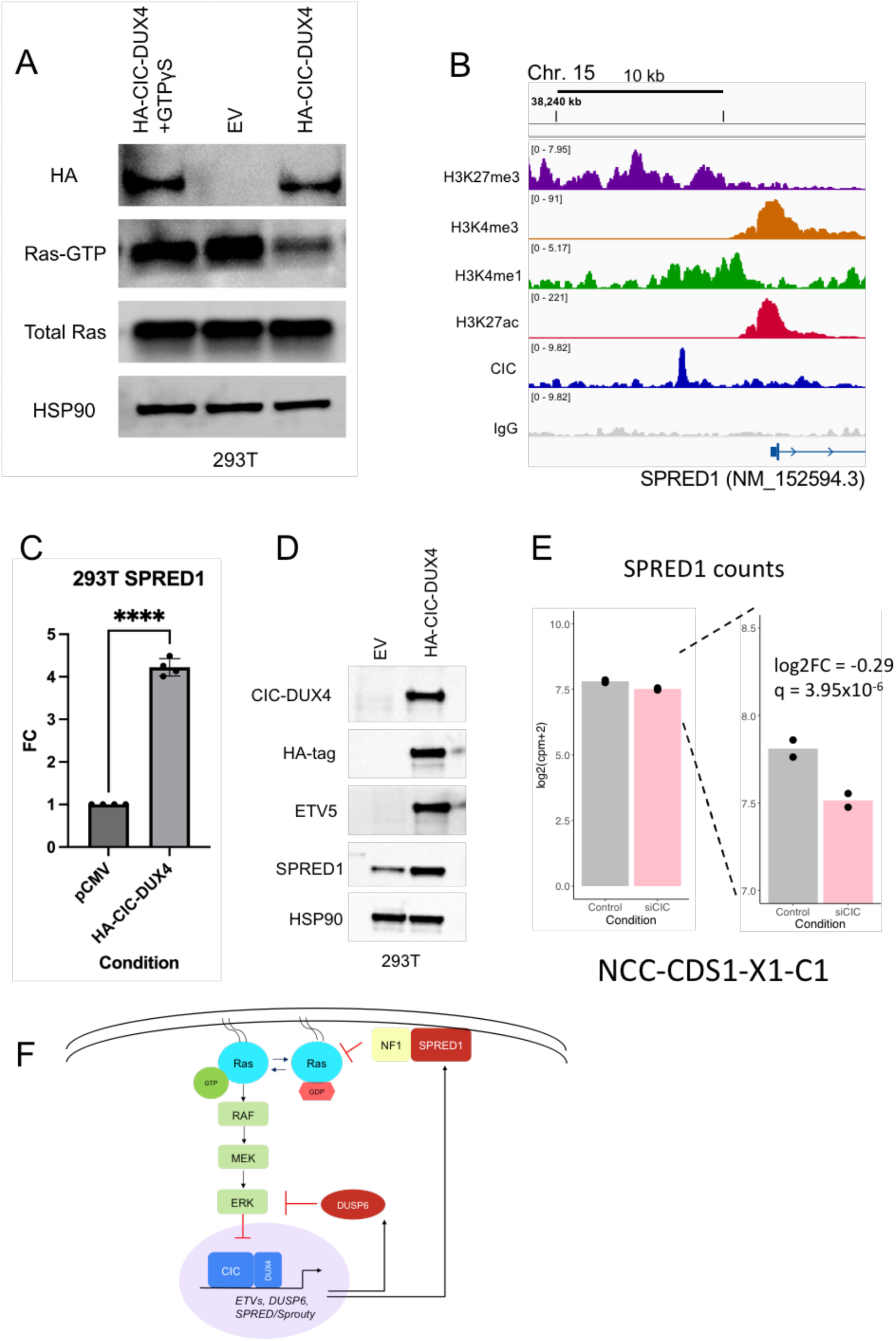
CIC-DUX4 target genes negatively regulate the MAPK-RAS-ERK signaling pathway. A) Ras-GTP pulldown in 293T cells expressing EV, HA-tagged CIC-DUX4, or HA-tagged CIC-DUX4 +GTPγS (positive control). B) H3K27me3, H3K4me3, H3K4me1, H3K27ac, CIC, and IgG ChIP-seq tracts aligned at a SPRED1 regulatory element. The CIC-DUX4 peak was called by MACS2 as highly significant (-log10(q) = 15.75), but it was not included in the set of 392 high-confidence CIC-DUX4 (Figure 1A) peaks because it was not located within 1kb of a called H3K27ac or H3K4me3 peak. C) SPRED1 mRNA expression in 293T cells expressing pCMV control or HA-tagged CIC-DUX4 (FC=fold change, ****p<0.0001, calculated by student’s t-test, error bars represent standard error of mean). D) Immunoblot of CIC-DUX4, HA-tag, ETV5, SPRED1, and HSP90 control in 293T cells expressing EV or HA-tagged CIC-DUX4. E) Expression of SPRED1 mRNA in NCC-CDS1-X1-C1 cells from edgeR-normalized log(cpm) values. Similar fold-change was seen in NCC-CDS2-C1 cells, but the result was not significant. q-value derived from the exact test employed in edgeR. F) Schematic of CIC-DUX4 regulation of key target genes that silence the MAPK-Ras-ERK signaling cascade.

## Discussion

The CIC-DUX4 fusion oncoprotein constitutes a rare diagnostic entity that is associated with poor clinical outcomes for patients. Through our integrative ChIP-Seq and transcriptomic analyses using patient-derived cells, we reveal how the CIC-DUX4 oncoprotein associates with chromatin accessibility and transcriptional state. We find a significant fraction of CIC-DUX4 localizes to distal regulatory elements and intronic regions in CIC-DUX4 sarcoma cells. CIC-DUX4 binding within these regions correspond to poised and active sites of transcription, suggesting that CIC-DUX4 controls gene expression largely through *cis*-regulatory elements. Notably, we did not observe a direct link between CIC-DUX4 and H3K27me3, a mark of transcriptional repression, suggesting that the primary function of CIC-DUX4 is to activate target gene expression.

Our data also suggest that slight variations in the consensus wild-type CIC octameric sequence may retain CIC-DUX4 binding specificity. In particular we note that “TG” sequences within the TGNNTGNN motif may represent obligate recognition sites for CIC-DUX4 binding in human sarcoma. Two questions that arise from this observation are: 1) how conserved TGNNTGNN motifs are as CIC recognition sites in non-cancer and non-human species; and 2) what is the role of TGNNTGNN repeats and how do they influence CIC-DUX4 binding and target gene regulation. We speculate that CIC-DUX4 binding to TGNNTGNN repeat sites may provide an evolved biological mechanism that enables tight transcriptional regulation and titration of key target gene levels. For instance, titrating higher doses of CIC-DUX4 can potentially bind un-occupied TGNNTGNN DNA-motifs to increase target gene levels and *vice versa*. These studies our beyond the scope of our manuscript and should be the focus of future investigation.

We and others have previously defined the molecular mechanisms that sustain CIC-DUX4 expression. Specifically, CIC-DUX4 expression is regulated by MAPK-ERK flux, whereby active ERK binds CIC-DUX4 and leads to its nuclear export and degradation in the cytosol. In order to maintain its own expression, CIC-DUX4 transcriptionally upregulates the ERK specific phosphatase, *DUSP6.* DUSP6 silences ERK activity to maintain CIC-DUX4 expression. Through our analysis we now identify other negative regulators of the MAPK-ERK cascade that represent putative CIC-DUX4 target genes. These include multiple SPRED and SPROUTY family members that primarily feedback on proximal MAPK substrates including Ras to ensure tight regulatory control of the MAPK signaling cascade. Of these uncharacterized targets, *SPRED1* is regulated in a positive and negative manner upon CIC-DUX4 overexpression and depletion, respectively. SPRED1 is known to interact and localize with NF1 (Ras-GTPase) at the plasma membrane where it inhibits Ras activity. Thus, we hypothesize that CIC-DUX4 may upregulate SPRED1 to silence Ras and, in turn, MAPK flux via NF1 localization to the plasma membrane. These functional studies further indicate that CIC-DUX4 can regulate its own expression through multilevel negative regulation and feedback on the MAPK-Ras-ERK pathway.

## Materials and Methods

### Cell Lines and Culture

Cell lines were cultured as recommended by the American Type Culture Collection (ATCC). HEK293T (RRID:CVCL_HA71) cells were purchased from ATCC. NCC_CDS1_X1_C1 (RRID:CVCL_YL54) and NCC_CDS2_C1 (RRID:CVCL_YL70) cells were generated as patient-derived cell lines and validated previously (17,18). All cell lines were authenticated using STR profiling. NCC-CDS1-X1-C1 and NCC-CDS2-C1 cells were grown RPMI 1640 media supplemented with 10% FBS, 100 IU/mL penicillin and 100 mg/mL streptomycin, and HEK293T cells were grown DMEM media supplemented with 10% FBS, 100 IU/mL penicillin and 100 mg/mL streptomycin. All cell lines were maintained at 37C in a humidified atmosphere at 5% CO2.

### RNA Extraction and Sequencing

Isolation and purification of RNA were performed using an RNeasy Mini Kit (QIAGEN), including an on-column DNase digest. RNA quality was evaluated using an Agilent TapeStation 4150 (Agilent).

Library preparation was performed by Novogene through poly-A selection and mechanical fragmentation followed with the NEBNext Ultra II RNA Library Prep for Illumina kit and sequenced on a Novaseq 6000 (paired-end 150bp reads) platform.

### Quantitative RT-PCR

Isolation and purification of RNA were performed using an RNeasy Mini Kit (QIAGEN). Total RNA (1000 ng) was used in a reverse transcriptase reaction with the SensiFAST cDNA Synthesis Kit (Bioline). Quantitative PCR included 3 replicates per cDNA sample. Human CIC (also targeting CIC-DUX4), SPRED1, SPRED2, SPRY4, ETV5, and GAPDH were amplified with Taqman gene expression assays (ThermoFisher Scientific). Expression data were acquired using a StepOnePlus Real-Time PCR System (Applied Biosystems). Expression of each target was calculated using the 2^−ΔΔCt^ method, and mRNA levels are expressed relative to GAPDH.

### Western Blot

All immunoblots represent at least 2 independent experiments. Adherent cells were washed and lysed with RIPA buffer supplemented with proteinase and phosphatase inhibitors. Proteins were separated by SDS-PAGE, transferred to nitrocellulose membranes, and blotted with antibodies recognizing HSP90 (Cell Signaling 4874, RRID:AB_2121214), CIC (Origene AP50924PU-N,), ETV5 (Cell Signaling 16274), SPRED1 (Cell Signaling 94063, RRID:AB_2800221), HA-Tag (Cell Signaling 3724).

### Gene Knockdown and Overexpression Studies

ON-TARGET plus Non-Targeting Pool (# D-001810-10-20), and CIC siRNA (#L-015185-01-0005) were obtained from GE Dharmacon and transfection was performed with Lipofectamine RNAiMax transfection reagent (Thermo Fisher Scientific).

The HA-CIC-DUX4 construct was a kind gift from Takuro Nakamura, Tokyo, Japan (7). HA-CIC-DUX4 was expressed in HEK293T cells using Fugene 6, according to the manufacturer’s protocol.

### Active RAS Pulldown Assay

The RAS GST-RBD activation kit was obtained from Cytoskeleton (Denver, CO, USA; cat #BK008). The protocol was performed following the manufacturer’s instructions. Lysis buffer for the Ras-GTP pulldown was 50 mM Tris (ph 7.5), 10 mM MgCl2, 0.5 M NaCl, and 2% Igepal. Snap-freezing of lysates using liquid nitrogen baths was performed directly after lysis. CIC-DUX4 was expressed in HEK293T cells and compared to empty vector (negative control) and GTPγS (positive control).

### ChIP-seq Bioinformatic Analysis

CIC-DUX4 immunoprecipitation was performed with NCC_CDS_X1 cells using the SimpleCHIP Enzymatic Chromatin IP Kit (Cell Signaling Technology) with IgG (Cell Signaling Technology) and CIC (Acris-Origene), H3K27ac (Active Motif #39133), H3K4me1 (Abcam #ab8895), H3K4me3 (Cell Signaling Technology #9751S, RRID:AB_2616028), and H3K27me3 (Cell Signaling Technology #9733, RRID:AB_2616029) antibodies in accordance with the manufacturer’s protocol. Paired-end 150-bp (PE150) sequencing on an Novaseq 6000 platform was then performed.

FASTQ files were checked for quality using fastqc (30) and multiqc, (31)and no major issues were noted. For sequence alignment, the human RefSeq reference genome GRCh38.p14 was downloaded from NCBI and indexed using bwa (32). BWA-MEM (Sniffles, RRID:SCR_017619) was used to align all samples to the reference genome, and samtools (33) was used to convert the resulting SAM files to BAM files. Picard (34) was used to name-sort and mark duplicates before position-sorting to index BAM files. Samtools (SAMTOOLS, RRID:SCR_002105) was used to generate flagstat reports for duplicate-marked, position-sorted BAM files. Picard-calculated duplication rates varied between 12% and 25% depending on the sample. Flagstat reports indicated mapping rates of 97-99+% and proper pairing rates of 94-98+% for all samples.

To call peaks, we used MACS2 (24) through a Docker image (fooliu/macs2). Replicates were combined by providing both replicate BAM files for the IP or control condition at the same time. For all conditions, we specified the -g hs, -f BAMPE, --keep_dup 1, and -q 0.1 parameters. For the CIC, H3K4me3, and H3K27ac conditions, we additionally enabled -- call-summits. For the H3K4me1 and H3K27me3 conditions we enabled --broad and omitted --call-summits. Called peaks were further filtered for more stringent q values using the awk command.

To determine a set of high-confidence CIC-DUX4 peaks, we used the bedtools (BEDTools, RRID:SCR_006646) (35) window command to identify CIC peaks with q < 0.001 within 1kb of a H3K27ac or H3K4me3 peaks, as these are marks for transcriptional activation. We then used bedtools to remove any peaks overlapping with known problematic regions using hg38-blacklist.v2 from the Boyle Lab (https://github.com/Boyle-Lab/Blacklist/blob/master/lists/). Peaks were further subset to just those on chromosomes (as opposed to patches or other sequence types), yielding the 392 high-confidence CIC-DUX4 peaks.

For visualization in IGV (36), BAM replicates were combined using samtools merge and indexed using Picard. deeptools (37) was used to generate bigWig files from these merged BAM files with the parameters --binSize 25, -e, --ignoreDuplicates, -- normalizeUsing RPGC, --effectiveGenomeSize 2913022398, and -bl with the same blacklist used above. deeptools was further used to generate QC plots and heatmap/profile plots of ChIP signal at specified locations: either high-confidence CIC-DUX4 peaks or H3K27me3 peaks (q < 0.001, not overlapping with blacklisted regions). R/RStudio (R version 4.2.2, R Core Team 2022) (38–40) were used throughout for simple plotting and data cleaning, with those scripts available at https://github.com/cuylerluck/CICDUX4_ChIP_RNA_seq. HOMER (41) was used for peak annotation using the hg38 reference genome.

For DNA motif analysis, 500bp sequences centered around the middle coordinate of high-confidence CIC-DUX4 peaks were extracted from the GRCh38.p14 sequence using bedtools. Since a small number of peaks were distinct summits within the same general peak region, yielding separate peaks with identical genomic coordinates, only unique genomic regions were included to avoid overrepresentation of identical sequences in the motif analysis. This resulted in 374 unique 500bp sequences being used. We used the browser version of the MEME Suite (42) for analysis, using MEME-ChIP (v5.5.2) in classic mode with default inputs and human DNA – HOCOMOCO Human (V11 CORE) as the motif database.

### ChIP-seq Track Visualization

IGV (2.15.2) was used for visualization of BAM and bigWig files with GRCh38/hg38 set as the genome. When BAM files from ChIP-seq data were visualized, the “Filter duplicate reads” option was enabled. This was also true of visualizing bigWig files, but not relevant as they were generated with the deeptools option --ignoreDuplicates included. Visualization of specific DNA motifs was done using the “Find Motif” tool embedded in IGV.

### Octamer Variant Bioinformatic Analysis

Full documentation of this analysis is available at https://github.com/cuylerluck/CICDUX4_ChIP_RNA_seq. Briefly, the 392 high-confidence CIC-DUX4 peaks were first reduced to 374 unique genomic regions based on the chromosome, start coordinate, and end coordinate of the peaks. As described in the ChIP-seq analysis section, this is because a small number of peaks were distinct summits within the same general peak region, and we did not want to double-count any identical peak sequences for motif analysis. We then used bedtools to extract the FASTA sequences within these 374 unique genomic peak regions from the reference genome used in the ChIP-seq analysis, GRCh38.p14.

Peak sequences were then loaded into R using RStudio for further analysis. We designed a function that given a peak sequence, generates every possible subsequence of length 8 (overlapping allowed), and returns both the raw frequency of each unique 8-mer and the proportion of all total 8-mers that a given 8-mer comprised (i.e. instances of 8-mer divided by total number of 8-mers). Given the results of our MEME-ChIP analysis, we then simplified the proportion data to only retain 8-mers that followed the pattern TGNNTGNN (or the reverse complement, NNCANNCA), where N is any base. We calculated the average frequency of this subset of 8-mers across all sequences and averaged these values between 8-mers that were reverse complements of each other. This was then visualized using standard ggplot2.

Because the average frequency of a given 8-mer could be large for two different reasons, either that it occurs often in many sequences or very frequently in a small subset, we then took the raw counts of 8-mers and similarly subsetted them to TGNNTGNN (or NNCANNCA) 8-mers and merged reverse complement pairs. This data was visualized for all peak sequences and select peaks of interest using pheatmap. Information on what genomic element a given peak was annotated for was taken from HOMER annotations generated as described in the ChIP-seq analysis methods section.

### RNA-seq Bioinformatic Analysis

STAR (43) was first used to build a genome index for the *Homo sapiens* reference genome GRCh38.p13, which was downloaded from the ENSEMBL FTP server along with the matching GTF file. We used --sjdbOverhang 149 per the recommendation of max(ReadLength-1), given that our reads were 150bp. Next, STAR was used to align all samples to the reference genome while simultaneously generating gene counts using -- quantMode GeneCounts. All samples mapped well, with rates of uniquely mapped reads of at least 89%. We also used samtools to generate and index BAM files from SAM files. fastqc and multiqc were used to check raw fastq files for quality control, with no unexpected findings noted. In-house bash scripts using the grep command were used to check fastq files for specific sequences spanning the CIC-DUX4 breakpoint or wild-type CIC sequences. deeptools bamCoverage was used to generate bigWig files from the first replicate of each sample’s BAM file, and the --normalizeUsing CPM option was included to normalize values. IGV was used to visualize bigWig files.

Remaining analyses were performed in R using RStudio, and the script used is available at https://github.com/cuylerluck/CICDUX4_ChIP_RNA_seq. Of note, read counts generated by STAR were loaded and simplified to columns 1 and 2 since the library preparation was unstranded. A standard edgeR (edgeR, RRID:SCR_012802) pipeline using the qCML method and the exact test was run for each cell line individually. Genes significantly downregulated after siCIC treatment in at least one cell line were written to a text file and used as input into DAVID (20) (default settings, Homo sapiens) and PANTHER (21) (default settings, biological process, Homo sapiens, GO Ontology database DOI: 10.5281/zenodo.7709866 released 2023-03-06). From these outputs, selected GO terms were plotted. Routine R scripting was used for all other analyses.

## Author Contributions

NJT, CL, and NS designed and performed the experiments and analyzed the data. RKP and ZK analyzed the data and provided critical revisions on the manuscript. RAO directed the project, analyzed experiments, and wrote the manuscript.

## Acknowledgements

RAO was supported by grants from the National Cancer Institute (K08CA222625 and R37CA255453) and the Children’s Cancer Research Fund. NJT was supported by a Research Fellowship from the UCSF School of Medicine. CL is supported by the UCSF Discovery Fellows program. NS is supported by an NCI Diversity Supplement (R37CA255453-03S1).

