## Supplementary data - 4 figures for "Mapping chromatin state and transcriptional response in CIC-DUX4 undifferentiated round cell sarcoma"

The supplementary data file contains 4 figures.

### Supplemental Figure 1

A

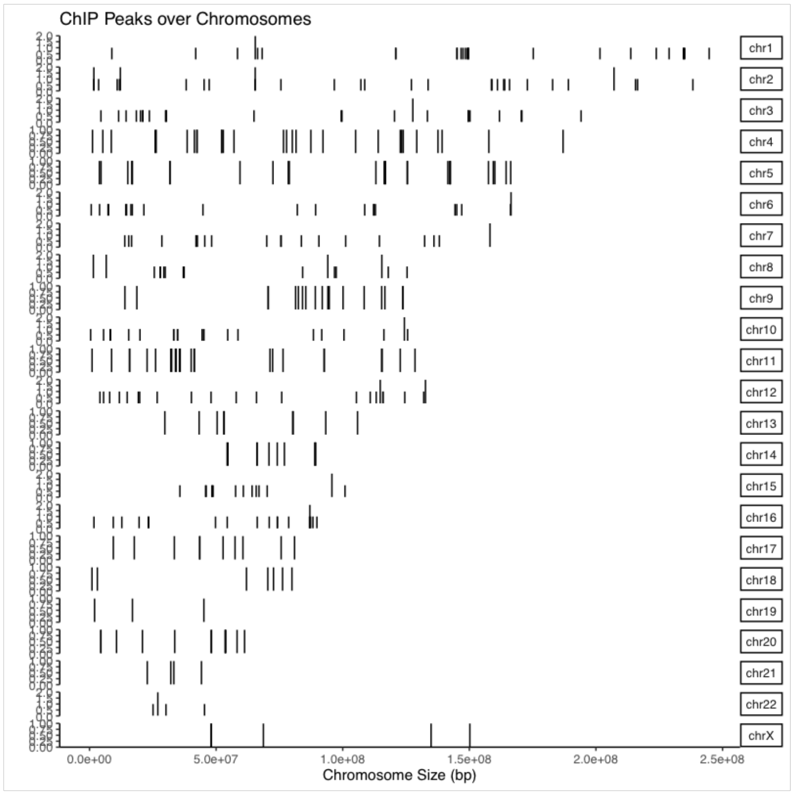


B

CIC-DUX4 peaks annotated as being at promoters (n = 12)

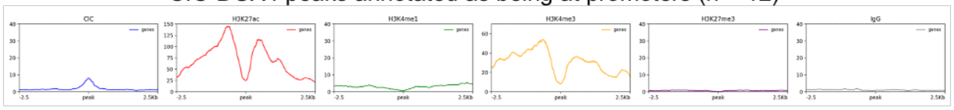


C

CIC-DUX4 peaks annotated as being intergenic (n = 174)

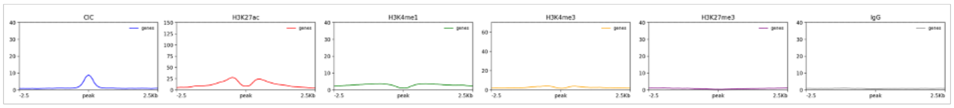


**Supplemental Figure 1: Chromosomal localization of CIC-DUX4 ChIP-seq peaks.**

A) Chromosome locations of high-confidence CIC-DUX4 peaks. B) deeptools profile plots using data analyzed in Figure 1C, subset into peaks annotated by HOMER as being located in promoters or intergenic regions.

Supplemental Figure 2

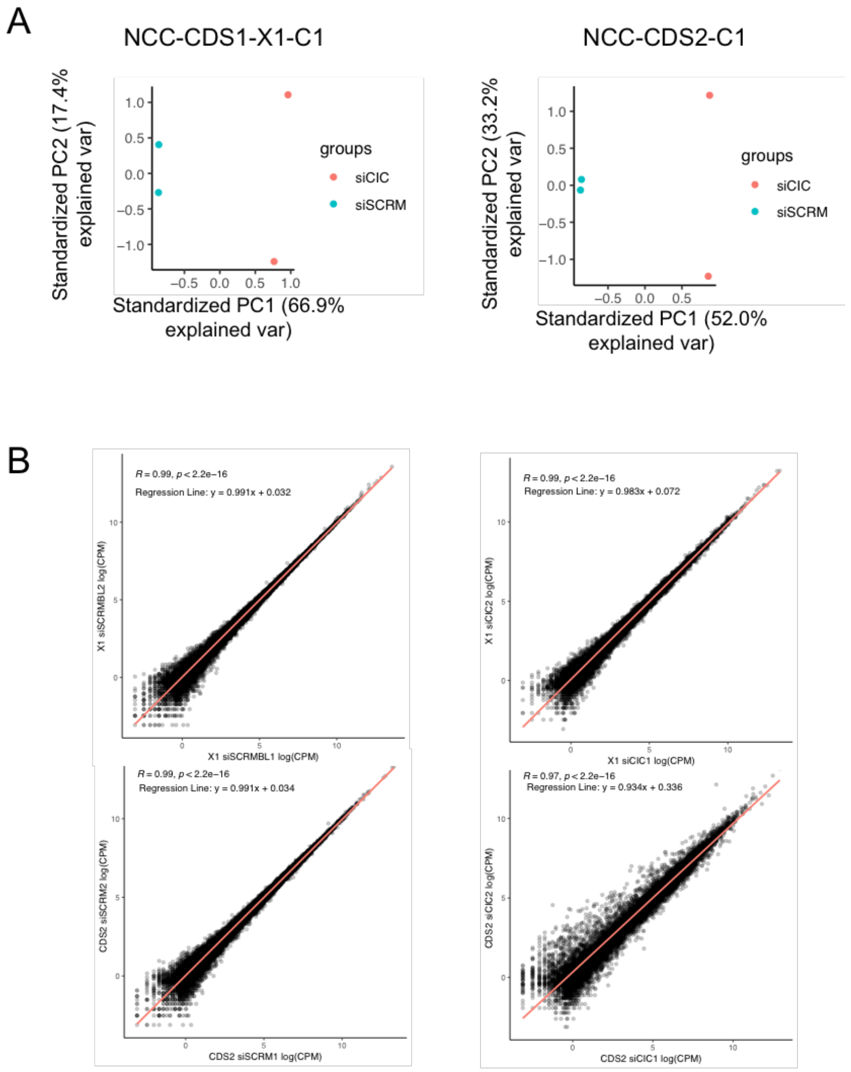


**Supplemental Figure 2: CIC-DUX4 regulated transcriptional changes in patient-derived cell lines.** A) PCA plots using normalized log(cpm) values retrieved following edgeR analysis for samples in each cell line. Values were centered and scaled as part of the fast.prcomp function call. B) Correlations between replicates for each experimental condition using normalized log(cpm) values retrieved following edgeR analysis. Statistics displayed include the pearson coefficient (R) and the best fit line determined by a y~x linear regression.

Supplemental Figure 3

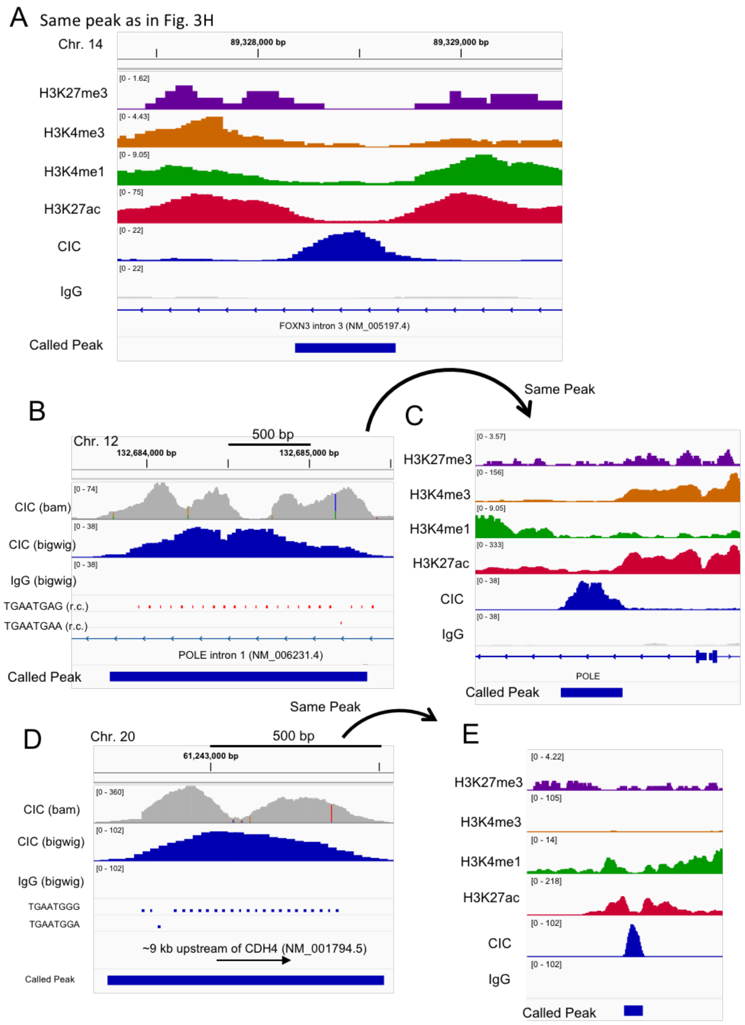


**Supplemental Figure 3: Non-canonical CIC binding motifs associate with CIC-DUX4 ChIP-seq peaks**. A) Same peak as in Figure 3H visualized in IGV aligned with designated histone marks tracks. B) IGV visualization of the CIC-DUX4 peak in *POLE* intron 1, including TGNNTGNN-variant locations. (r.c.) = reverse complement). C) Visualization of the same peak as in B, aligned with tracks for histone marks shown. D) IGV visualization of the CIC-DUX4 peak upstream of *CDH4*, including TGNNTGNN-variant locations. E) Visualization of the same peak as in D, aligned with tracks for histone marks shown.

Supplementary Figure 4

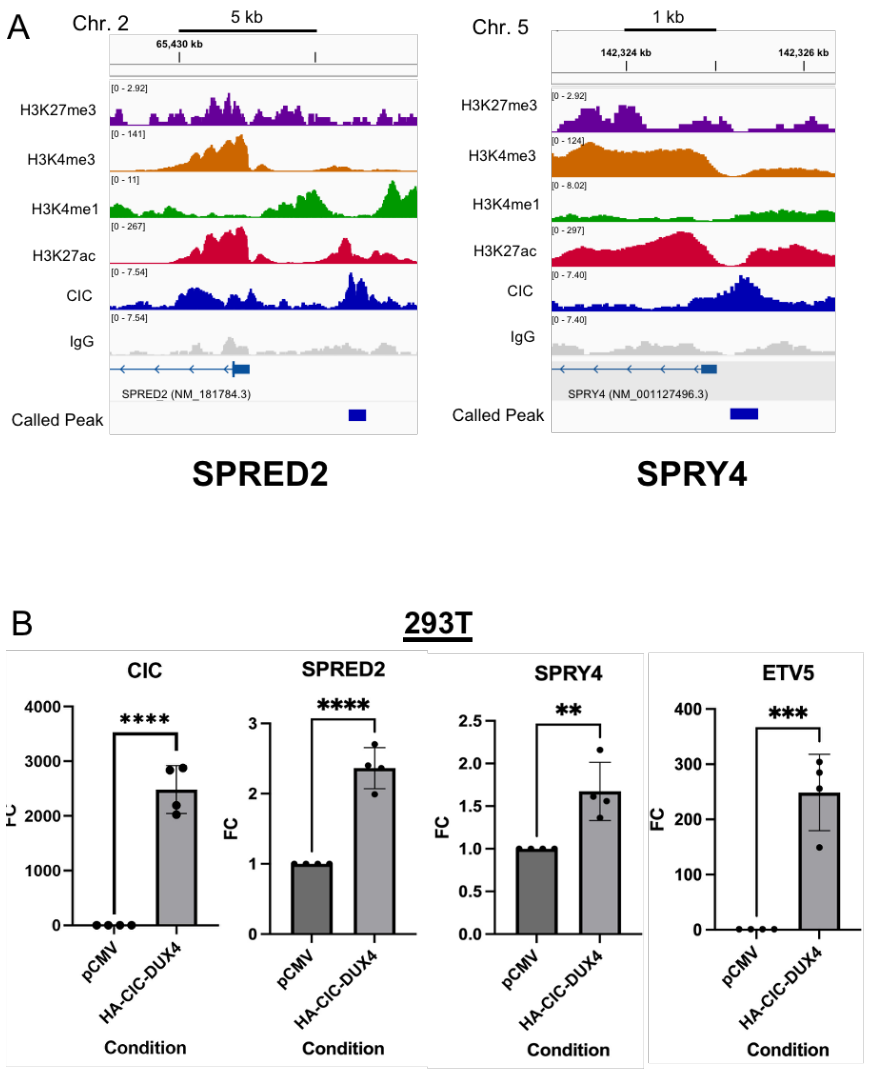


**Supplementary Figure 4. CIC-DUX4 peaks localize to negative regulators of MAPK-RAS signaling.**

A) H3K27me3, H3K4me3, H3K4me1, H3K27ac, CIC, and IgG ChIP-seq tracts aligned at putative *SPRED2* (A) and *SPRY4* (B) regulatory elements. Visualized by IGV. B) *CIC, SPRED2, SPRY4, ETV5* mRNA expression in 293T cells expressing pCMV control or HA-tagged CIC-DUX4 (FC=fold change, **p<0.01, ***p<0.001, ****p<0.0001, calculated by student's t-test, error bars represent standard error of mean).
